# Heterogeneous Firing Responses Predict Diverse Couplings to Presynaptic Activity in mice Layer V Pyramidal Neurons

**DOI:** 10.1101/091587

**Authors:** Yann Zerlaut, Alain Destexhe

## Abstract

In this study, we present a theoretical framework combining experimental characterizations and analytical calculus to capture the firing rate input-output properties of single neurons in the fluctuation-driven regime. Our framework consists of a two-step procedure to treat independently how the dendritic input translates into somatic fluctuation variables, and how the latter determine action potential firing. We use this framework to investigate the functional impact of the heterogeneity in firing responses found experimentally in young mice layer V pyramidal cells. We first design and calibrate *in vitro* a simplified morphological model of layer V pyramidal neurons with a dendritic tree following Rall's branching rule. Then, we propose an analytical derivation for the membrane potential fluctuations at the soma as a function of the properties of the synaptic input in dendrites. This mathematical description allows us to easily emulate various forms of synaptic input: either balanced, unbalanced, synchronized, purely proximal or purely distal synaptic activity. We find that those different forms of input activity lead to various impact on the membrane potential fluctuations properties, thus raising the possibility that individual neurons will differentially couple to specific forms of activity as a result of their different firing response. We indeed found such a heterogeneous coupling between synaptic input and firing response for all types of presynaptic activity. This heterogeneity can be explained by different levels of cellular excitability in the case of the balanced, unbalanced, synchronized and purely distal activity. A notable exception appears for proximal dendritic inputs: increasing the input level can either promote firing response in some cells, or suppress it in some other cells whatever their individual excitability. This behavior can be explained by different sensitivities to the speed of the fluctuations, which was previously associated to different levels of sodium channel inactivation and density. Because local network connectivity rather targets proximal dendrites, our results suggest that this aspect of biophysical heterogeneity might be relevant to neocortical processing by controlling how individual neurons couple to local network activity.

## Introduction

The specific activation of subpopulations within neocortical networks appears to be the core mechanism for the cortical representation of sensory features. The details of how such specific activations happen are therefore key questions in systems neuroscience. As a primary source for specific activation, the neocortex is characterized by some degree of specific circuitry: neurons differ in their afferent connectivity. A classic example can be found in the primary visual cortex, layer IV simple cells specifically sample their input from ON and OFF cells in the thalamic nucleus [1,2]. Additionally, neocortical neurons also vary in their electrophysiological properties: for example, heterogeneous levels in the action potential threshold are routinely measured *in vivo* [3–5]. Thus, an emerging refinement is that the sensitivity of a neuron to a given feature do not only results from its *stimulus specificity* (e.g. orientation selectivity as a result of a specific afferent circuitry), but from the combination of its *stimulus specificity* and its *biophysical specificity*. Two somato-sensory cortex studies illustrates this point precisely. In Crochet et al. [3], during active touch, the spiking probability of a neuron (its sensitivity to whisker touch) follows from the combination of the reached level of synaptically-driven membrane potential deflection (its *stimulus-specificity* resulting from afferent circuitry, as quantified by post-synaptic *reversal potentials*) and its threshold for action potential triggering (its *biophysical specificity*). A similar result was found in the study of Yang et al. [5] for texture recognition, where the combination of those two quantities was shown to predict choice-related spiking. Those results therefore suggest that heterogeneity in the biophysical properties of neocortical neurons might have an impact on their functional role during sensory processing.

#### Author summary

Neocortical processing of sensory input relies on the specific activation of subpopulations within the cortical network. Though specific circuitry is thought to be the primary mechanism underlying this functional principle, we explore here a putative complementary mechanism: whether diverse biophysical features in single neurons contribute to such differential activation. In a previous study, we reported that, in young mice visual cortex, individual neurons differ not only in their excitability but also in their sensitivities to the properties of the membrane potential fluctuations. In the present work, we analyze how this heterogeneity is translated into diverse input-output properties in the context of low synchrony population dynamics. As expected, we found that individual neurons couple differentially to specific form of presynaptic activity (emulating afferent stimuli of various properties) mostly because of their differences in excitability. However, we also found that the response to proximal dendritic input was controlled by the sensitivity to the speed of the fluctuations (which can be linked to various levels of density of sodium channels and sodium inactivation). Our study thus proposes a novel functional impact of biophysical heterogeneity: because of their various firing responses to fluctuations, individual neurons will differentially couple to local network activity.

In the present work, we further investigate the interaction between *stimulus specificity* and *biophysical specificity* in the light of the variability in the biophysical features reported in our previous study [6], namely that single neurons in juvenile mice cortex not only vary in their excitability (linked to the action potential threshold) but also in their sensitivity to the properties of the membrane potential fluctuations. Our previous communication introduced those new dimensions in the *biophysical specificity*, we aim here at understanding their functional impact. To this purpose, we implemented various stimuli onto layer V pyramidal cells (we varied the properties of presynaptic activity in the fluctuation-driven regime), and we investigated whether individual neurons would differentially respond to those inputs as a result of their various firing rate responses [6] (their various *biophysical specificities*).

## Results

### Single cell computation in the fluctuation-driven regime: input variables and output quantity

Our study investigates the properties of single cell computation in the regime of low synchrony population dynamics [7,8] (the analogous, at the network level, of the fluctuation-driven regime at the cellular level) and aims at describing effects mediated by slow population dynamics (*T* ≥20-50ms). In this context, the cellular input-output function of a neocortical neuron corresponds to the function that maps the presynaptic variables to the spiking probability of the neuron, which is often called the *transfer function* of the neuron.

The framework of our approach to the transfer function is illustrated in Figure 1A. Our cellular model has five presynaptic variables: four of them are presynaptic firing rates (stationary release probabilities at the synapses) as those constitute the primary input variables in this rate-based paradigm. To investigate their differential contribution, the proximal and distal parts of the dendritic trees have been treated separately and each of them has two presynaptic rates corresponding to the excitatory and inhibitory input (hence four rate variables: 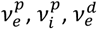 and 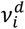). The main motivation for this separation is to distinguish between two types of projections onto neocortical pyramidal neurons: synaptic inputs from the local network are thought to be more proximal while the distal apical tuft receives input from more distant cortical areas and thalamic locations [9]. Additionally, a global synchrony variable has been introduced for presynaptic events. This reproduces the effect of multi-innervation of a cell by its presynaptic afferent and, more importantly, the effect of pairwise correlations associated to neocortical dynamics [10]. The synchrony degree in the presynaptic activity has been suggested to vary with stimulus statistics in the primary visual cortex [11,12] what motivates its introduction as a separate variable in our model.

### A two-step approach to determine the cellular input-output function

As illustrated on Figure **1**A, we introduce a two-step procedure to determine the input-output function of a single cell in the fluctuation-driven regime. The idea behind this two-step approach relies on the fact that the action potentials are initiated at the axon initial segment [13] (i.e. electrotonically close to the soma) so that the fluctuations at the soma could determine the firing probability uniquely (see Discussion for the approach’s limitations).

We thus split the relation from presynaptic quantities (the input) to the spiking probability (the output) into the following successive steps. The first step consists in evaluating how dendritic integration will shape the membrane potential fluctuations at the soma for given values of the presynaptic input variables. The fluctuations are quantified by their mean *μ*_*V*_, their amplitude *σ*_*V*_ (given by the standard deviation of the fluctuations) and the fluctuations speed *τ*_*V*_ (given by the autocorrelation time of the fluctuations). This first step will be performed analytically as the passive properties and simplified morphology of the dendritic model enables a mathematical treatment of this question. The second step consists in determining how the somatic fluctuations (*μ*_*V*_, *σ*_*V*_, *τ*_*V*_) are translated into action potential output. This last step is computed thanks to a fitted function directly constrained by experiments. Indeed, by analyzing the spiking response of pyramidal neurons recorded *in vitro* as a function of these somatic fluctuations (*μ*_*V*_, *σ*_*V*_, *τ*_*V*_), we previously found that a parametric function could reliably describe the relation between the fluctuations properties and the firing rate output in individual neurons [6]. Thus, because the experiments were performed as a function of these somatic variables (*μ*_*V*_, *σ*_*V*_, *τ*_*V*_), the same experimental data obtained previously can be used in the present framework.

Within this framework, one individual cell (indexed by *k*) is therefore described by two functions: 1) a dendritic integration function 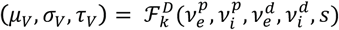 that accounts for synaptic integration up to the somatic *V*_*m*_ fluctuations and 2) a parametric function 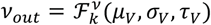 that translates the somatic *V*_*m*_ fluctuations into a spiking output. The final input-output function 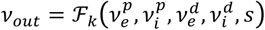 of cell *k* is thus the result of the composition: 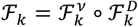 (see Figure **1**A). From our previous study, we benefit from a set of firing response functions 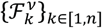 of n=30 cells. To complete the framework, we now need to associate a dendritic morphology for each of this cell to be able to calculate the associated set of dendritic integration functions 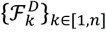. This is the focus of the next two sections. We first present our dendritic morphology model and then derive an association rule from input resistance to dendritic morphology based on *in vitro* recordings of somatic input impedance.

**Figure 1.**
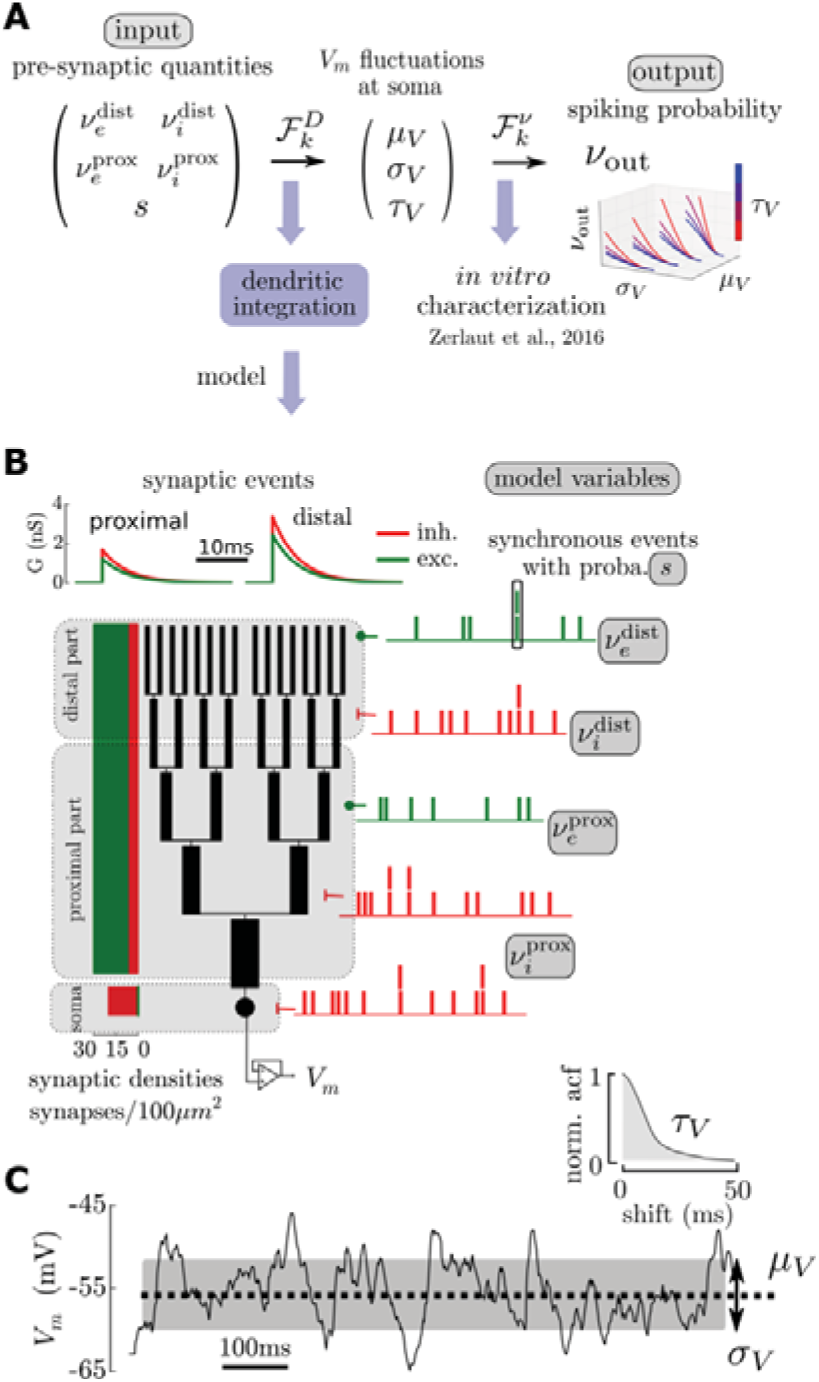
A theoretical framework for single cell computation in the fluctuation-driven regime: cellular model and two-step approach for the input-output function. (A) Theoretical paradigm: to get the input-output function of a single cell, we split the relation from presynaptic quantities (the input) to the spiking probability (the output) into two steps. 1) passive dendritic integration shapes the membrane potential at the soma and 2) how those fluctuations are translated into spikes is captured by a firing response function determined *in vitro*. (B) Theoretical model for dendritic integration. A single cell is made of a lumped impedance somatic compartment and a dendritic tree. The dendritic tree is composed of B branches (here B=5), the branching is symmetric and follow Rall's 3/2 rule for the branch diameters. Synapses are then spread all over the membrane according to physiological synaptic densities. We define 3 domains: a somatic and proximal domain as well as a distal domain, excitatory and inhibitory synaptic input can vary independently in those domains. An additional variable: synaptic synchrony controls the degree of coincident synaptic inputs. (C) A given presynaptic stimulation (here 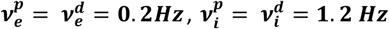 and 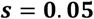) creates membrane potential fluctuations at the soma characterized by their mean ***μ***_***V***_, their amplitude ***σ***_***V***_ and their autocorrelation time ***τ***_***V***_.

### A simplified morphological model for dendritic integration

The morphology of our theoretical model is a lumped impedance somatic compartment in parallel with a dendritic arborescence of symmetric branching following Rall's 3/2 branching rule (see Figure **1**B and Methods). This morphology is of course a very reductive description of pyramidal cells: it does not discriminate between the distinct apical trunk and the very dense basal arborescence. Also, branching in pyramidal cell morphologies have been shown to deviate from Rall's 3/2 branching rule. Nonetheless this simplified model contains the important ingredient for our study: the fact that the transfer impedance to the soma of a synaptic input will strongly depend on its location on the dendritic tree. Indeed, as observed experimentally [14], distal events will be more low-pass filtered than proximal events in this model.

We spread synapses onto this morphology according to physiological densities [15] and describe synaptic events as transient permeability changes of ion-selective channels (see Methods). We arbitrarily separate the dendritic tree into two domains: a proximal and a distal domain (delimited by their distance to the soma, see Figure **2**B). The distal part was taken as the last eighth of the dendritic tree to reproduce the large electronic distance to the soma characterizing distal synapses [14]. Following experimental evidences [14], we set a higher synaptic efficacy for distal synapses. The synaptic parameters take physiological values [16] and can be found on Table 1. The passive parameters as well as the individual morphologies are estimated in the next section.

**Figure 2.**
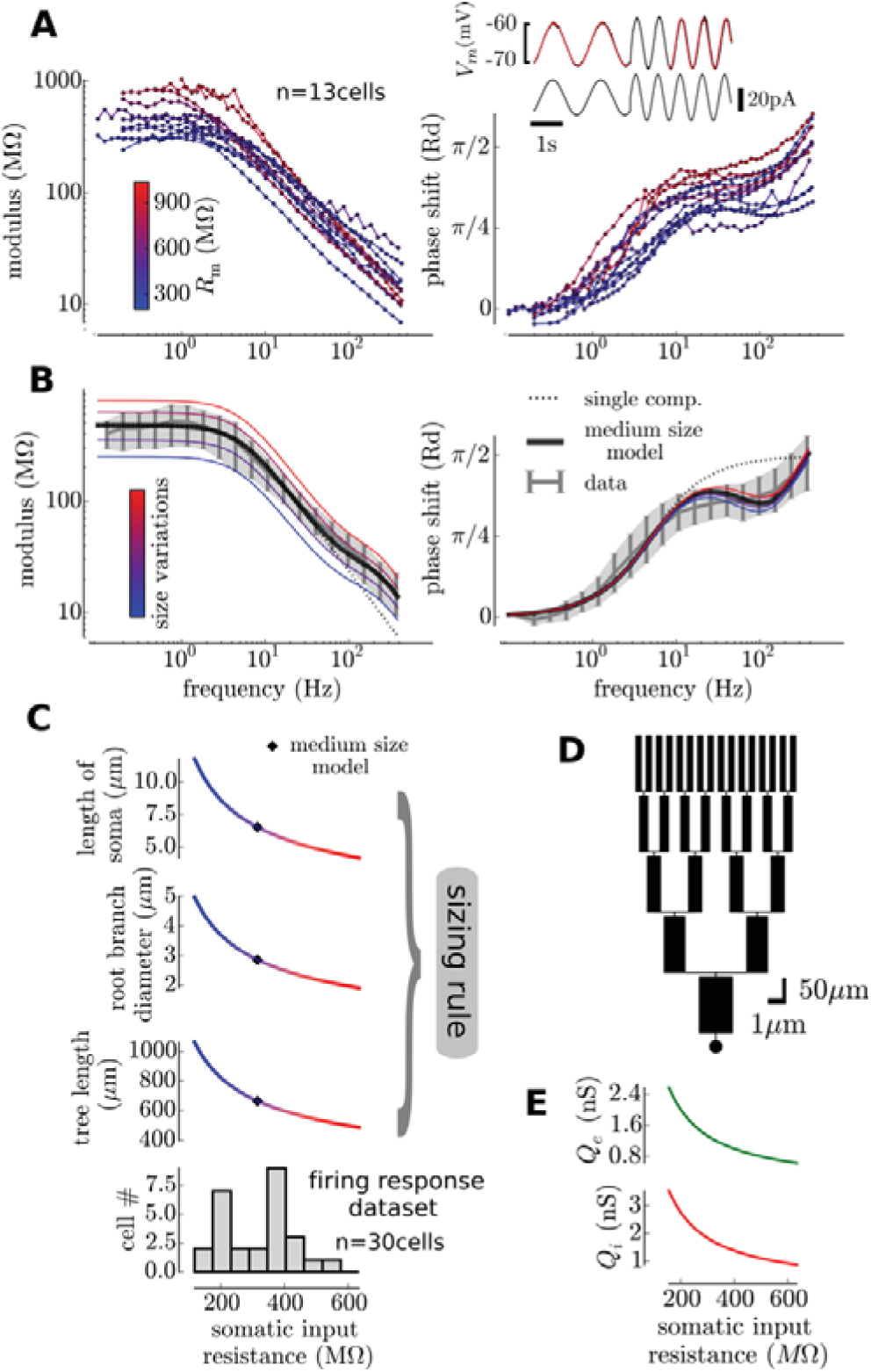
Calibrating the model on *in vitro* measurements: the simplified model and its size variations provides an approximation for the somatic input impedance of pyramidal cells and its heterogeneity over the recorded population. (A) Input impedance (left: modulus and right: phase shift) measured at the soma in intracellular recordings with sinewave protocols in current-clamp (inset). The color code indicates the input resistance and is likely to result from size variations of individual cells. (B) A medium size model accounts for the average data and varying the size of the dendritic tree and soma (according to the sizing rule shown in C) partially reproduces the variability in the individual measurements. Large cells (blue) have a lower modulus and a lower phase shift while small cells (red) have both a higher modulus and phase shift. (C) We obtain a map between input resistance and size of the morphological model. (D) Representation of the medium-size model. (E) Additionally the synaptic weights are rescaled with respect to the cell's somatic input resistance. Because the mean transfer resistance to soma is linked to the input resistance, this rescaling insures that the mean synaptic efficacy at soma is the same for all cells.

**Table 1.** Model parameters

| Parameters | Name | Symbol | Value | Unit |
| --- | --- | --- | --- | --- |
| Passive |  |  |  |  |
| | Leak conductance density | $G_L$ | 325 | $\mu\text{S}/\text{cm}^2$ |
| | Intracellular resistivity | $R_i$ | 30 | $\Omega.\text{cm}$ |
| | Specific capacitance | $C_m$ | 1.05 | $\mu\text{F}/\text{cm}^2$ |
| | Leak reversal potential | $E_L$ | -65 | mV |
| Synaptic |  |  |  |  |
| | Inhibitory reversal potential | $E_i$ | -80 | mV |
| | Excitatory reversal potential | $E_e$ | 0 | mV |
| | Somatic inhibitory density | $\mathcal{D}_i^{\text{soma}}$ | 20 | Synapses/(100$\mu\text{m}^2$) |
| | Somatic excitatory density | $\mathcal{D}_e^{\text{soma}}$ | 0 | Synapses/(100$\mu\text{m}^2$) |
| | Dendritic inhibitory density | $\mathcal{D}_i^{\text{tree}}$ | 6 | Synapses/(100$\mu\text{m}^2$) |
| | Dendritic excitatory density | $\mathcal{D}_e^{\text{tree}}$ | 30 | Synapses/(100$\mu\text{m}^2$) |
| | Proximal inhibitory weight | $Q_i^{0,p}$ | 1.0 | nS |
| | Proximal excitatory weight | $Q_e^{0,p}$ | 0.7 | nS |
| | Distal inhibitory weight | $Q_i^{0,d}$ | 1.5 | nS |
| | Distal excitatory weight | $Q_e^{0,d}$ | 1.05 | nS |
| | Inhibitory decay time | $\tau_i$ | 5 | ms |
| | Excitatory decay time | $\tau_e$ | 5 | ms |
| Mean Morphology |  |  |  |  |
| | Soma length | $l_s$ | 5.0 | $\mu\text{m}$ |
| | Soma diameter | $d_s$ | 15.0 | $\mu\text{m}$ |
| | Root branch diameter | $d_t$ | 2.25 | $\mu\text{m}$ |
| | Tree length | $l_t$ | 550.0 | $\mu\text{m}$ |
|  | Branch number | B | 5 |  |
| | Proximal tree fraction | $f_{\text{prox}}$ | 7/8 | |

### Estimating the passive properties and individual morphologies of layer V pyramidal cells

We will use the firing response functions of the cells of our previous study [6] (the “firing response dataset”). For each of this cell, we therefore need an estimate of the parameters of the dendritic model (described above, i.e. passive properties and morphology parameters). The purpose of this section is thus to derive an association rule from the input resistance (a quantity that we have for all cells of the “firing response dataset”, see Figure **2**C) to the parameters of the dendritic model. We based this estimate on the comparison between the dendritic model behavior and the properties of the somatic input impedance in layer V pyramidal neurons (n=13 cells, measured with intracellular recordings *in* vitro).

The key property on which this characterization relies is the fact that the input impedance at the soma cannot be accounted for only by the isopotential somatic compartment (i.e. a RC circuit). The input impedance shows the contribution of the dendritic tree in parallel to the soma [17]. Indeed, both the modulus and the phase of the input impedance show deviations from the RC circuit impedance (see the comparison in Figure **2**B): see for example the exponent of the power law scaling of the modulus (-1 exponent for the single compartment and ∼-0.7 for pyramidal cells) or the decreased phase shift around 100Hz. As this behavior is the consequence of the electrotonic profile along the dendritic tree, we used it to estimate the parameters of our simplified dendritic model.

We first average all data (shown on Figure **2**A) to obtain a mean input impedance (shown on Figure **2**B) representative of a mean cellular behavior. We then performed a minimization procedure to obtain both the passive properties and the morphology corresponding to this average behavior (see Methods). The obtained passive properties were compatible with standard values, e.g. the resulting specific capacitance was 1.05 *μ*F/cm^2^, close to the commonly accepted 1 *μ* F/cm^2^ value, thus suggesting that the procedure could capture the physiological parameters of pyramidal cells, see Table 1 for the other parameters. Most importantly, the surface area was physiologically realistic, so that when using synaptic densities, we obtain an accurate number of synapses (see below). A representation of this mean morphology can be seen on Figure **2**D.

Pyramidal cells show a great variability in input impedance, for example their input resistance almost spans one order of magnitude (both in the present n=13 cells, see the low frequency modulus values in Figure **2**A, as well as in the firing response dataset, see bottom in Figure **2**C). We found that varying the size of the morphological model within a given range around the mean morphological model could partially reproduce the observed variability in the input impedance profiles (see Figure **2**B). Size variations corresponds to a linear comodulation of the 1) tree length *L*_*t*_, 2) the diameter of the root branch *D*_*t*_ and 3) the length of the somatic compartment *L*_*S*_ (see Figure **2**C for the range of their variations). On Figure **2**A, the cells have been colored as a function of their input resistance while on Figure **2**B, we vary the size of the size of the morphological model. Large cells (blue, low input resistance) tend to have a lower input resistance and phase shift than the small cells (red, high input resistance). Note that this simplistic account of morphological variations only very partially describes the observed behavior in pyramidal cells. In particular, 1) it strongly underestimates the variations of phase shifts at medium and high frequencies (f>20Hz) and 2) the relationship between size and impedance modulus at high frequencies (f>100Hz) is poorly captured. Those discrepancies are likely to be due to the details of dendritic arborescence that are not captured by the strong constraints of our dendritic model (symmetric branching, diameter rules, number of branches, etc…). Despite those discrepancies, size variations in our morphological model constitute a reasonable first approximation to account for cellular variety within the layer V pyramidal cell population.

This characterization, combined with the analytical tractability of the model (see Methods) allowed us to construct a map between input resistance at the soma and size of the morphological model (the passive properties are set as identical, the one fitted on the mean impedance behavior). Thus, for each neuron in our previous "firing response dataset", because we have its input resistance at the soma, we can associate a given morphology. The association rule is shown in Figure **2**C.

We now check what is the number of synapses obtained from the combination of our fitted morphologies with the physiological synaptic densities. We found a number of synapses of 3953 ± 1748 (mean and standard deviation across the n=30 cells) with a ratio of excitatory to inhibitory numbers of synapses of 4.5 ± 0.1. The fact that those numbers fall within the physiological range constitutes a validation of our approach (the morphology estimate through input impedance profile characterization).

### An analytical approximation for the properties of the membrane potential fluctuations at the soma

We now want to translate the five variables of the model in terms of membrane potential fluctuations properties at the soma (*μ*_*V*_, *σ*_*V*_, *τ*_*V*_), i.e. determining the function 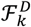 for each cell *k* depending on its dendritic parameters. This constitutes the first step to obtain the final input-output function of individual cells (see Figure **1**A).

**Figure 3.**
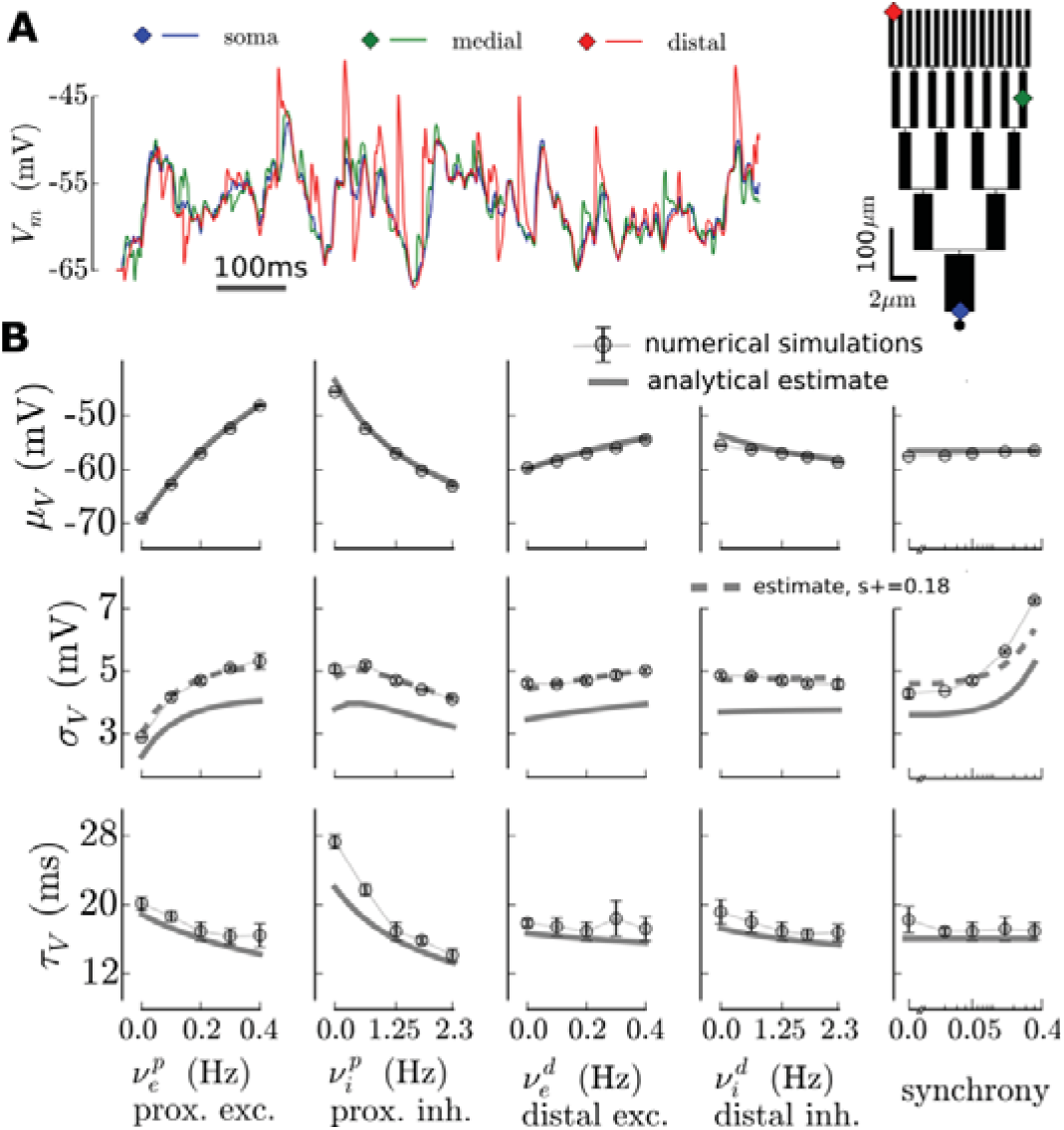
Accuracy of the analytical estimate for the properties of the membrane potential fluctuations at the soma: comparison between numerical simulations and the analytical approximation. Shown for the medium size model of Figure 2D. (A) In the numerical simulation, we explicitly simulate the whole dendritic arborescence, we show the membrane potential variations for the three locations shown on the left. (B) Properties of the membrane potential fluctuations (mean ***μ***_***V***_, standard deviation ***σ***_***V***_ and autocorrelation time ***τ***_***V***_) for different configuration of presynaptic activity: analytical predictions and output from numerical simulations in NEURON. In each column, one variable is varied while the other variables are fixed to the mean configuration value corresponding to 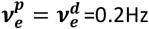, 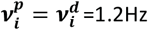 and s=0.05. In the ***σ***_***V***_ plots (middle panels, dashed gray lines), we added the prediction of the analytical estimate after a +0.18 correction for the synchrony (found with a Newton method).

Investigating dendritic integration for detailed morphological structures is made difficult by the fact that this has to be done numerically with a relatively high spatial and temporal discretization. In the fluctuation-driven regime, one also needs to sample over long times (T ≫ *τ*_*V*_ ∼ 20ms) to obtain the statistical properties of the somatic *V*_*m*_ resulting from dendritic integration. In addition, this study relies on n=30 different morphologies and we will explore a five dimensional parameter space (the five variable of our model). Under those conditions, if performed numerically, the computational cost of such a study is clearly prohibitive. We briefly describe here, why, in our simplified model, an analytical treatment is possible and thus renders this investigation feasible (see details in the Methods and in the Supplementary Material). The key ingredient is the ability to reduce the dendritic tree to an equivalent cylinder [17], we only adapted this reduction to the changes in membrane permeability associated to the high conductance state [18]. Two approximations underlie our estimation: 1) the driving force during an individual synaptic event is fixed to the level resulting from the mean bombardment [19] and 2) the effect at the soma of a synaptic event at a distance *x* in a branch of generation *b*, corresponds to the 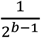 fraction of the post-synaptic response to the stimulation made of synchronous events at distance *x* in all the 2^*b*-1^ branches of the generation *b*. Luckily, the combination of those approximation is a favorable situation. Indeed, hypothesis 1) overestimates the size of post-synaptic events (because the driving force is not fixed, it diminishes during the PSP time course) while hypothesis 2) underestimates the size of post-synaptic events (because of the 2^*b−1*^ − 1 synchronous events in neighboring branches, the membrane conductance is higher than in the case of a single event, consequently neighboring events have a shunting effect that artificially decreases the response). In addition, both of those approximation are likely to hold when single events are of low amplitude compared to the amplitude of the massive synaptic bombardment (see e.g. Kuhn et al. [19] for the validity of the first hypothesis).

In Figure **3**, we compare the analytical approximation to the output of numerical simulations performed with the *NEURON* software [20]. We varied the five variables of the model around a mean synaptic bombardment configuration (see next section). Some discrepancies between the approximation and the simulations appeared, in particular one can see a ∼1mV shift in the standard deviation *σ*_*V*_ of the fluctuation (meaning that single events are underestimated in the analytical treatment, so that hypothesis 2 is the most problematic one). Because the synchrony controls the amplitude of the fluctuations (Figure 3B and next section), the analytical estimate could therefore be seen as an accurate estimate, modulo a shift in the synchrony (see Figure 3B, an increase of 0.18 in the synchrony corrects for the ∼1mV shift in *σ*_*V*_). Importantly, the trend in the variations of the fluctuations as a function of the model variables is globally kept between the analytical estimate and the numerical simulations. This relatively good agreement therefore shows that our analytical estimate is a valid tool to study dendritic integration in the fluctuation-driven regime.

### Properties of the fluctuations for different types of presynaptic activity

We now implement various types of presynaptic activity and investigate the properties of the resulting membrane potential fluctuations at the soma. In addition, we represent the variations of the somatic input conductance (relative to the leak input conductance) because, as it is routinely measured in intracellular studies *in vivo*, this quantity allows a comparison between the model and experimentally observed activity levels. On Figure **4**, we present those different protocols, on the left (panel **A**), one can see how the five variables of the model are comodulated for each protocol (color coded, see bottom legend) and on the right (panel **B**), one can see the resulting properties of the membrane potential fluctuations. We present those results only for the medium-size model, but it was calculated for the morphologies associated to all cells. The variability introduced by the various morphologies is shown in Figure S2 and we found that the qualitative behavior discussed in this section was preserved in all cells. We first introduced a baseline of presynaptic input corresponding to a low level of network activity: 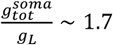, compared to ∼3-4 in activated states, reviewed in [18]. This baseline activity is a mix of proximal and distal activity with a low degree of synchrony (*S*=0.05). Similarly to Kuhn et al. [19], the inhibitory activity is adjusted to obtain a balance of the *V*_*m*_ fluctuations at -55mV. The firing values of this baseline level are very low 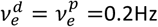 for the excitation and 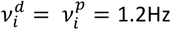 for the inhibition) in accordance with the sparse activity characterizing mammalian neocortical dynamics [3,10]. On top of this non-specific background activity, we will now add a specific stimulation. We consider four types of presynaptic stimulations:

- *Unbalanced activity increase*. We define unbalanced activity as a stimulation that brings the mean membrane potential above -55mV corresponding to the previously defined balance. The stimulation corresponds to an increase of the excitatory synaptic activity (still running within a very sparse range of activity, 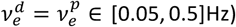 with an increasing inhibitory activity adjusted to linearly disrupt the balance between -55mV and -52mV (see Figure **4**). The synchrony is kept constant and the activity indifferentially raises in the proximal and distal part. This increase of total activity raises the input conductance ratio close to four. In this moderate range, the variations of the amplitude of the fluctuations *σ*_*V*_ remains a monotonic increase (unlike the non-monotonic variations found in the single-compartment study of Kuhn et al. [19] and the case of a proximal stimulation, see below), the fluctuations gets approximately twice faster (the normalized autocorrelation time 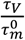 decays from 100% to 50%) and, of course (by design), the mean depolarization has a linear increase of 3mV.
- *Proximal activity increase*. To emulate purely proximal activity, we fix the distal presynaptic firing frequencies (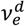 and 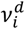) as well as the synchrony to their baseline levels. To remain in a sparse activity level, we increase the proximal excitatory activity from the baseline level to 1.7Hz and we adjust proximal and somatic inhibitory activity to keep the balance at the soma. This would nonetheless correspond to large network activity level, as can be seen from the input conductance ratio (that raises up to 8). This situation gives results comparable to the single-compartment study of Kuhn et al. [19]. The amplitude of the fluctuations has a non-monotonic profile and the autocorrelation time strongly decreases. A notable difference is that, even if we investigated high activity levels, the autocorrelation time does not go to zero and the amplitude of the fluctuations has only a moderate decrease. This discrepancy is due to 1) the choice of non-negligible synaptic time constants compared to the membrane time constants (here *τ*_syn_ =5ms and 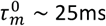, then 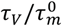 would saturate at 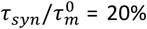 and 2) the strong shunting effects observed in the single compartment case is here attenuated because the synaptic input is distributed.
- *Distal activity increase*. For distal activity, we keep the proximal presynaptic frequencies (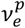 and 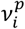) as well as the synchrony to their baseline levels. We increase the distal excitatory activity 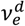 from the baseline level to a moderate level: 0.7Hz. The distal inhibitory frequency 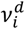 is again adjusted to keep the balance at the soma. Here, we get a different picture than in the proximal case, the increase in activity leads to negligible increase of the somatic input conductance as expected from electrotonically distant input [21]. Also, the decrease of the speed of the fluctuations is much attenuated. The reason for this phenomenon is that only the distal part has a high conductance, consequently post-synaptic events are strongly low-pass filtered by the proximal part of the arborescence before reaching the soma. Here, the amplitude of the fluctuations strongly increases as a function of the input and do not show the non-monotonic relation found for proximal input. This is explained by the combination of the fact that 1) distal events are of higher amplitude and 2) the shunting of post-synaptic events is much reduced due to the relatively narrow localization of the synaptic conductances.
- *Increase in synaptic synchrony.* Finally, we emulate an increase in the presynaptic synchrony. Here, all synaptic frequencies are kept constant with respect to the baseline level and we simply increase the probability of coincident events for each synaptic spike train. Because there is no change in synaptic activity, this stimulation does not affect the input conductance ratio, neither the mean membrane potential or the speed of the fluctuations. However, presynaptic synchrony strongly affects the amplitude of the fluctuations in a near linear manner.

**Figure 4.**
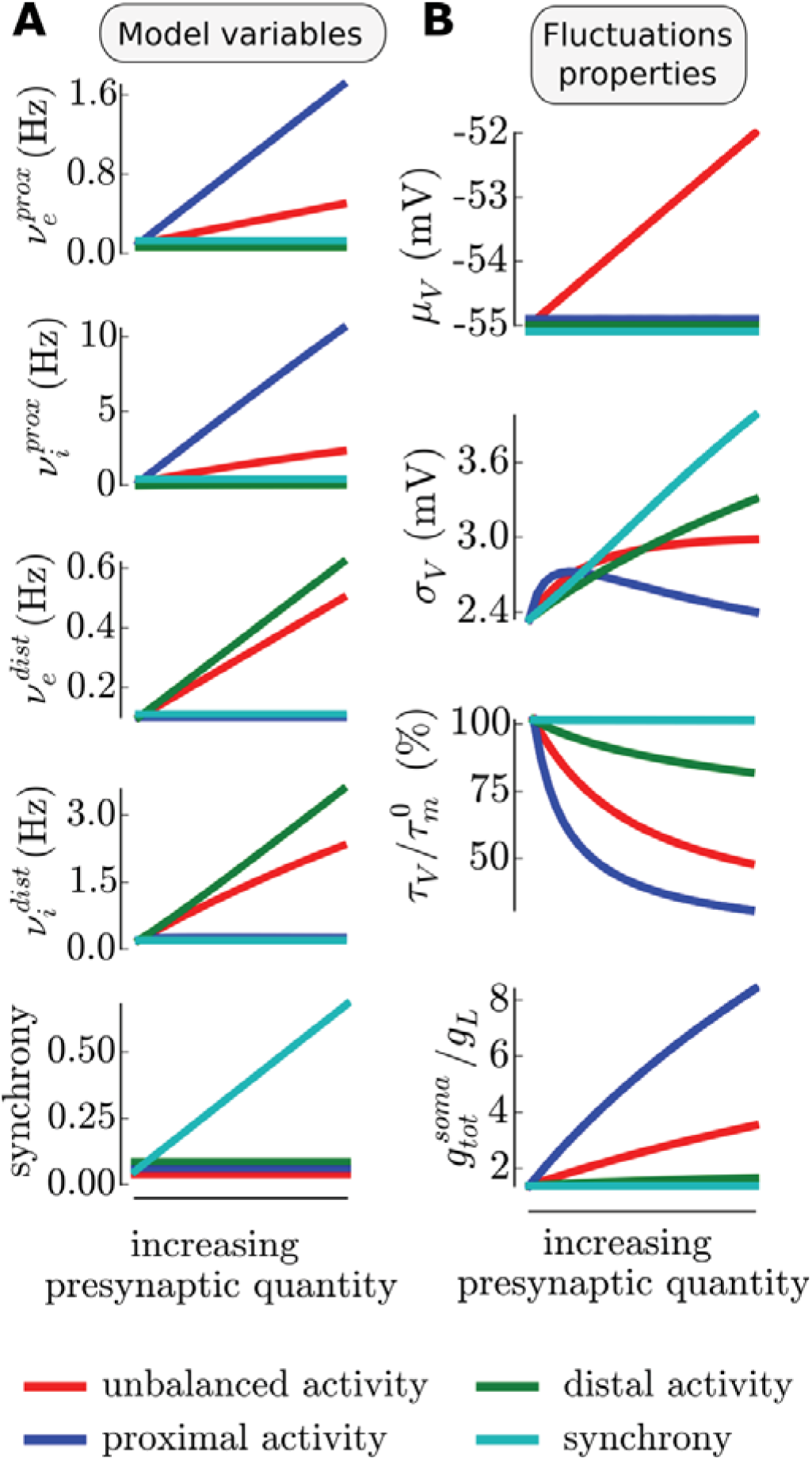
Properties of the membrane potential fluctuations for various types of presynaptic activity: either unbalanced (red), purely proximal (blue), purely distal (green), synchronized (cyan). A common baseline configuration of balanced proximal and distal activity at low rate gives rise to baseline fluctuations properties, on top of this, the increase of a given type of presynaptic activity corresponds to a given comodulations of the 5 model variables. (A) Comodulations of the model variables to achieve varying levels of the different types of activity. (B) Membrane potential fluctuations properties (mean ***μ***_***V***_, standard deviation ***σ***_***V***_ and autocorrelation time ***τ***_***V***_) and somatic input conductance at the soma for the different protocols. Shown for the medium-size model, see Supplementary Material for the variability introduced by variations in cell morphologies.

Note, that in addition to the sparse activity constraints or the balance constraints, the criteria for the ranges of the model variables was chosen to have the fluctuations in the same domain. For example, we investigated a lower activity range for the distal part (variations of *ν*^*dist*^) than for the proximal part (variations of *ν*^*prox*^) to avoid an explosion of *σ*_*V*_, the range for the synchrony increase followed the same criteria.

### Heterogeneous firing responses induce diverse coupling to presynaptic activity

For each one of the n=30 cells of our previous study [6], we have 1) a morphological model (see previous sections) and 2) a firing response function 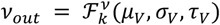. Thanks to the previous analytical approximation, we can translate the five model variables 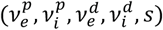 into the stationary fluctuations properties (*μ*_*V*_, *σ*_*V*_, *τ*_*V*_) that, in turn, the function 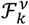 translate into a spiking probability. Thus, we finally get the full *input-output* function (within our theoretical framework) as illustrated on Figure **1**A.

We show on Figure **5** the response of four cells to the different types of presynaptic activity described in the previous section (those four cells were chosen as they were representative of different firing response behaviors, see Figure 5 and 6B in [6]). The input-output relationships show qualitative and quantitative differences, we briefly discuss them here and we perform a more rigorous analysis on the full dataset in the next section.

First, we can see that individual cells have a very different level of response to the baseline level of synaptic activity (initial response in Figure **5**). Cell 1 has a baseline at ∼ 10^−2^ Hz while Cell2 or Cell3 have response above 1Hz, i.e. two orders of magnitude above.

Importantly, those cells have different preferences for particular types of stimulations. Cell 1 responds more to unbalanced activity whereas Cell 2 and Cell 4 respond more to an increase in synchrony and Cell 3 responds preferentially to proximal activity (within this range). This is what we mean by *preferential coupling*: individual neurons will respond preferentially to a particular type of synaptic activity. An even more pronounced discrepancy appears for proximal activity: the response can be either increased (Cell 1 and Cell 3) or decreased (Cell 2 and Cell 4) with respect to the baseline level.

Given the relative invariance of the fluctuations properties for each cell (see previous section and Figure S2, despite the various morphologies, the same input creates the same fluctuations), those differences can only be attributed to the various firing responses of individual cells (the diversity in the 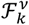 functions found experimentally [6]). We conclude that *heterogeneous firing responses induce diverse coupling to presynaptic activity*.

**Figure 5.**
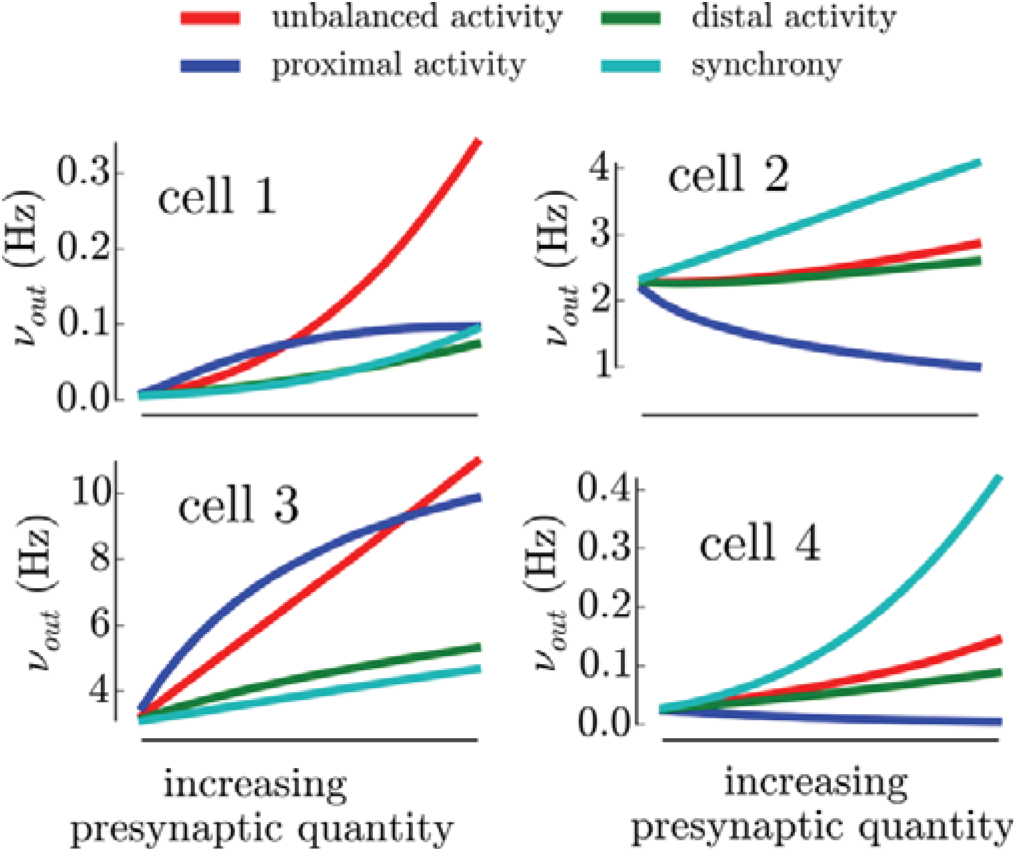
Examples of the firing response of 4 different cells for the various types of presynaptic activity (color-coded) shown in Figure 4. The abscissa “increasing synaptic quantity” corresponds to the comodulations of the model variables shown in Figure 4A (same color code). As an example, the response to a “proximal activity increase” (blue curve) corresponds to a linear increase of 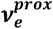 and 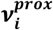 while keeping 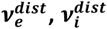 and *s* to their baseline level. See Figure 4A for the comodulations of the other types of increasing activity.

### Biophysical origin of the heterogeneous couplings to presynaptic activities

We now make this analysis more quantitative by computing the responses for all n=30 cells. We get their response to the baseline level *ν*_*bsl*_ and their mean response change for each stimulation type (the mean over the range of scanned presynaptic input): *δν*_*ubl*_ for the unbalanced activity, δ*ν*_*dist*_ for the proximal activity, δ*ν*_*dist*_ for the distal activity and δ*ν*_synch_ for an increased synchrony. We show the histogram of those values in the left column of Figure **6**.

We now investigate how the response of an individual cell relates to its *biophysical specificity*. It is defined here by four quantities [6] (see also Methods):

i. 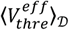: a measure of the cellular excitability.
ii. 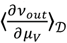: the sensitivity to the mean of the fluctuations, this quantifies how much a mean depolarization is translated into a change in firing rate.
iii. 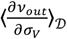: the sensitivity to the amplitude of the fluctuations, this quantifies how much an increase in the amplitude of the fluctuations is translated into a change in firing rate.
iv. 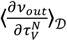: the sensitivity to the speed of the fluctuations, this quantifies how much a change in the speed of the fluctuations is translated into a change in firing rate.

We first analyze the response to baseline activity *ν*_*bsl*_. When log-scaled (Figure **6**A), the distribution is approximately normal and spans 2-3 orders of magnitude. This log-normal distribution of pyramidal cell firing rates during spontaneous activity seems to be a hallmark of mammalian neocortical dynamics (see e.g. [10] in human neocortex). We investigated what properties of the firing responses could explain this behavior, we therefore looked for correlations between our measures of the firing responses in the fluctuation-driven regime [6] and the baseline responses. Not surprisingly, we found a very strong linear correlation between the excitability 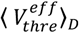 and the baseline response level, the other characteristics do not have an impact (Pearson correlations, see values in Figure **6**A). It should be stressed that presynaptic connections are homogeneous across cells in this model, those results therefore show that the typical log-normal distribution of firing rates could very naturally emerge as a result of the normal distribution observed in pyramidal cell's excitabilities [6], thus suggesting that no specific circuitry might be needed to explain this neocortical property.

**Figure 6.**
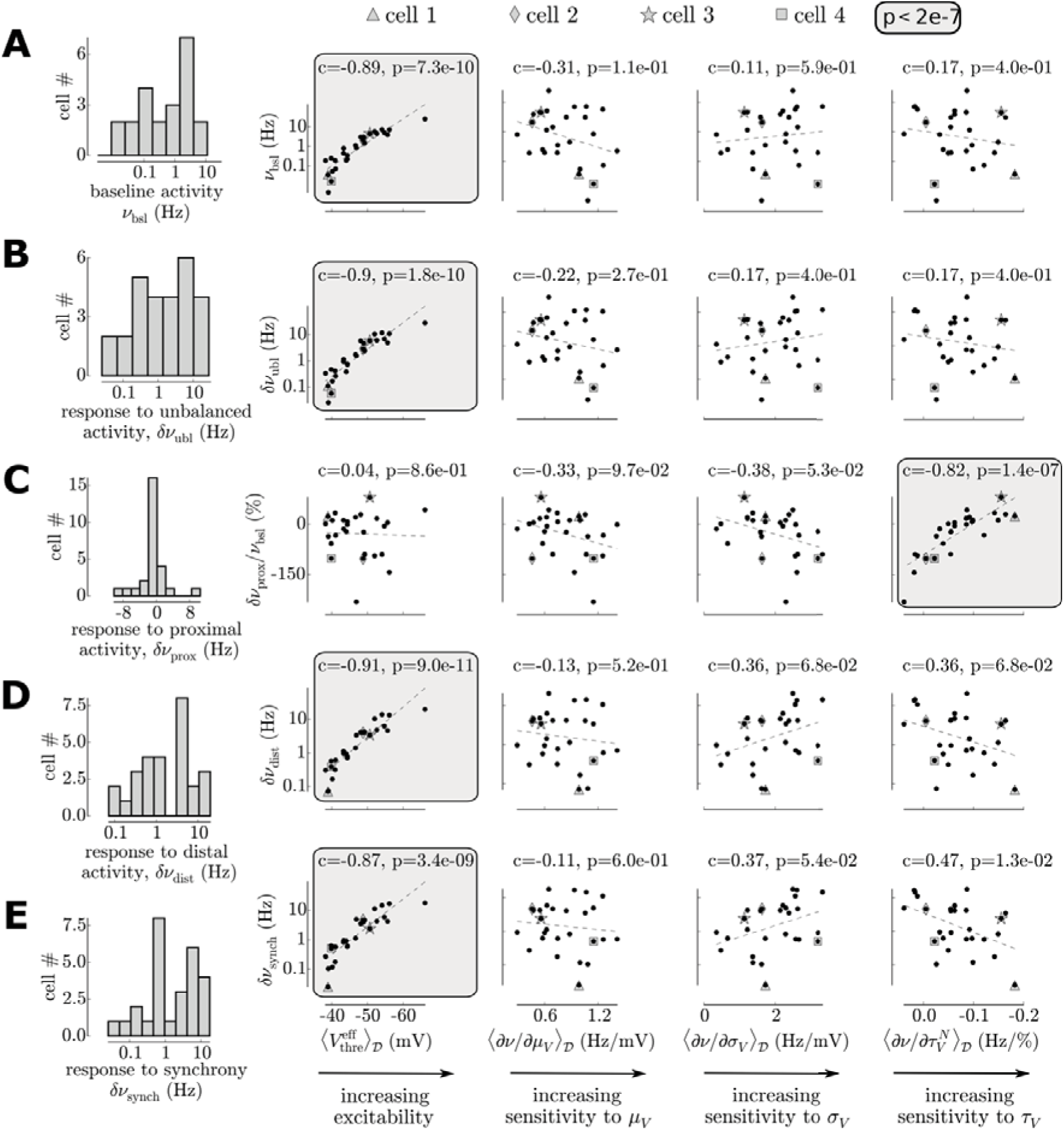
Diverse cellular responses to the various types of presynaptic activity and their link to the characteristics
of their firing response function. Note the logarithmic scale for the firing responses in B,C,D. (A) Diverse response to baseline stimulation. (B) Diverse response to unbalanced activity. (C) Diverse response to proximal activity. Note that because the response also show negative changes of firing rate, the data cannot be log-scaled. Instead, they have been rescaled by the baseline response (i.e. we show 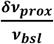). (D) Diverse response to distal activity. (E) Diverse response to a synchrony increase.

Despite the important differences in the fluctuations they create (see Figure **6**B), the responses over cells to unbalanced activity, distal activity and an increased synchrony share a very similar behavior. First, those stimuli produce systematically an increase in firing rate (n=30/30 cells). The firing increase again show a strong heterogeneity over cells, covering two orders of magnitude (see log y-axis on Figure **6**B,D,E). Again, this variability in responses was highly correlated with the excitability. Surprisingly, the response was not dependent on any other of the characteristics of the firing response. For example, because synchrony controls the standard deviation *σ*_*V*_, the variability observed during an increase in synchrony could have been linked to the to the sensitivity to the standard deviation 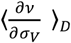, but this effect was not significant (see Pearson correlation in Figure **6**E). For those three protocols, none of the sensitivities to the fluctuations properties had a strong impact on the individual cellular responses (c<0.4 and p>0.01, Pearson correlations, see Figure **6**). This analysis therefore revealed that, for those type of synaptic activities, those properties of the firing response have negligible impact compared to the very strong effect of the variability in excitabilities (see Discussion).

The response to proximal activity also showed a great variability but with a qualitatively different behavior (Figure **6**C). Notably, firing could be suppressed or increased. This variability was independent of the excitability of the cells but was correlated with the sensitivity to the speed of the fluctuations 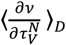. Indeed, the proximal stimulation implies a strong variations of the fluctuations speed (i.e. decreasing *τ*_*V*_, while keeping moderate variations of *σ*_*V*_ and, by design, a constant *μ*_*V*_) thus rendering the sensitivity to the fluctuation speed the critical quantity for this stimulation type. Those results therefore show that the response to proximal activity of an individual cell is controlled by its level of sensitivity to the speed of the fluctuations (see Discussion).

## Discussion

In the present work, we introduced an analytically tractable description of single cell computation in the fluctuation-driven regime for neurons with a dendritic structure. We introduced a two-step framework consisting of considering independently (1) how dendritic synaptic input translates into the properties of the somatic membrane potential fluctuations (*μ*_*V*_, *σ*_*V*_, *τ*_*V*_); (2) how these somatic variables translate into action potential output. We used this framework to investigate how the heterogeneity found experimentally in firing responses to fluctuating input shape the diverse inputoutput functions of neocortical pyramidal cells. Focusing on the regime of near asynchronous population dynamics, we emulated various types of presynaptic activity and found that those different types of synaptic stimulation correspond to various comodulations of the fluctuations properties. This property is what motivated to fully scan the three dimensional space (*μ*_*V*_, *σ*_*V*_, *τ*_*V*_) in our previous study [6] instead of the response to a given presynaptic input type that corresponds to an arbitrary comodulation of (*μ*_*V*_, *σ*_*V*_, *τ*_*V*_). Importantly, we found that, because of their different response to the same fluctuations, individual neurons would differentially couple to various types of synaptic inputs.

### A versatile theoretical framework for cellular computation in the fluctuation-driven regime

Despite the current weaknesses of our description (see next section), we believe that having an analytical model for dendritic integration in the fluctuation-driven regime is a useful tool for many problems in theoretical neuroscience. The main advantage of this model is that one can very naturally plug in physiological parameters (because surface area as well as transfer resistance to soma can take physiological values) while still allowing an analytical treatment (though see deviations of the approximations in Figure **3**). In the theoretical analysis of neural network dynamics, the literature is almost exclusively based on the reduction to the single-compartment (reviewed in [22]). Though being approximate, our framework thus opens the path toward a detailed mathematical analysis of recurrent network dynamics containing neurons with extended dendritic structures.

Additionally, It must be stressed that formulating the cell response with the somatic fluctuations properties (*μ*_*V*_, *σ*_*V*_, *τ*_*V*_) as an intermediate variable is very powerful because it allows one to apply the same measurements to various models. For example, with a single-compartment model it is easy to translate these variables into excitatory and inhibitory activities [19]. We showed here that it is also possible to obtain relations with synaptic inputs occurring in dendrites in a simplified morphological neuron model. The latter model uses the same measurements, so no experiments need to be redone. We could in principle also apply the same approach to more complex models and obtain more realistic transfer functions. This phenomenological two-step procedure thus offers a flexible complementary approach to the analytical approaches tackling the problem of the spiking behavior in presence of an extended dendritic structure [23–25].

### Limitations of the framework

The proposed theoretical framework for single-cell computation nonetheless suffers from several weaknesses. First, even if our description captures the various electrotonic distances associated to various synaptic locations (the crucial ingredient here to discriminate between proximal and distal inputs), the morphological model appears as a very poor description of layer V pyramidal cell. Deviations from the symmetric branching hypothesis and Rall's branching rule will have a significant impact on dendritic integration in the *fluctuation-driven* regime. To investigate those effects within the framework proposed in our study, one could benefit from the large body of theoretical work on the derivation of Green’s function for arbitrary branched passive dendritic trees [26–29]. Another important limitation of our description lies in the absence active mechanisms in dendrites [30]. It is therefore a question how much those mechanisms could affect the picture provided in our study. Preliminary numerical analysis performed in presence of NMDA and Ca^2+^ currents [31] (see Figure S4 and Figure S5), showed that, provided excitation balances inhibition (a situation where NMDA channels keeps a relatively low level of stationary activation) and provided synchrony do not reach a too high level (unlike for s ≥ 0.4, where excitatory events almost systematic lead to NMDA spikes), the qualitative behavior of the cellular input-output function remains unaffected. Thus, even if the lack of dendritic mechanisms underestimates the coupling values reported here (cells are less excitable in the passive setting, leading to attenuated spiking), the absence of qualitative differences suggest that the cell-to-cell variability reported in this study would be poorly affected. Finally, our description assumes a unique correspondence between average presynaptic quantities, somatic fluctuations and output spiking. The compartmentalized nature of active dendritic integration is very likely to break this hypothesis: the same average input could lead to very different output responses when targeting different functional subunits. We conclude that, given the complexity of synaptic integration in neocortical cells, the proposed approach only constitutes a very first approximation of the cellular input-output function. Quantifying its weaknesses and improving this picture should be the focus of future investigation.

### The key quantities of a single cell’s “biophysical specificity”: the excitability and the sensitivity to the speed of fluctuations

Very naturally, a key quantity to explain the various levels of neuronal responses is the cellular excitability. Indeed, the response to baseline activity, unbalanced activity, distal activity or an increase in synchrony is strongly correlated with cellular excitability.

More surprisingly, the sensitivity to the speed of the fluctuations also have a crucial impact on the response for one type of synaptic activity: proximally targeting synaptic input. In our previous communication [6], theoretical modeling suggested that a high sensitivity to the speed of the fluctuations was enable by a high level of sodium inactivation (as only fast fluctuations allow to deinactivate sodium channels) and a high density of sodium channels (as it corresponds to a sharp spike initiation mechanism that enables to extract fast fluctuating input, reviewed in [32]). The present analysis thus proposes an important functional role for those biophysical properties: controlling the coupling to proximally targeting activity. Additionally, the present analysis sheds light on the physiological relevance of the properties that we introduced in our previous study. In all four measures characterizing the cellular firing rate response function in the fluctuation-driven regime (shown at the bottom of Figure **6**), only two of them seem to be physiologically relevant: the excitability 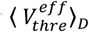 and the sensitivity to the speed of the fluctuations 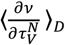. The reason for the lack of functional impact of the sensitivity to depolarizations 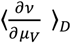 and to the sensitivity to amplitude of the fluctuations 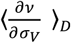 was that they were estimated independently of the excitability (after rescaling by the different excitability levels). While this separation was useful to isolate and evidence the contribution of different biophysical mechanisms to the firing rate response function [6], the firing response to a change in depolarization or an increase in amplitude of the fluctuations will be mainly led by the cell’s excitability level whereas the more subtle features of its firing response function (the sensitivities to *μ*_*V*_ and *σ*_*V*_) will have a negligible impact.

### Biophysical specificity might contribute to the diverse couplings to local network activity in neocortex

Finally, we speculate about a putative link between a recent observation and the present findings. The *in vivo* study in mice visual cortex of Okun and colleagues [4] reported a strong heterogeneity in the coupling between individual cell's responses and the locally recorded population activity. The authors explained those observations by a variability in the local recurrent connectivity and found that this diverse coupling did not seem to be explained by a variability in biophysical features (e.g. the coupling was independent from the action potential threshold, somehow a measure of the excitability). However, as local connectivity is thought to target more proximal regions such as the basal dendrites, our study proposes a biophysical mechanism that could also contribute to their observation. The diverse coupling to proximal activity was here explained by the sensitivity to the speed of the fluctuations, and similarly to their results, this coupling was found to be independent on the cellular excitability (see Figure 6B in Zerlaut et al. [6]). Part of their results could therefore be explained by this mechanism. Also, even if this electrophysiological heterogeneity disappears in mature phenotypes, the preferential coupling present in young animals could be amplified by long term plasticity to form this strongly coupled local network. Future work could therefore address this hypothesis by combining recordings of population activity with a subtle and functionally-relevant analysis of single cell properties.

## Methods

### Ethics Statement

Experiments were performed at Unité de Neurosciences, Information et Complexité, Gif sur Yvette, France. Experimental procedures with animals were performed following the instructions of the European Council Directive 2010 86/609/EEC and its French transposition (Décret 2013/118).

### Morphological model

The morphology of our theoretical model is the following (depicted in Figure 1B): it is made of an isopotential somatic compartment (i.e. a leaky RC circuit) in parallel with a dendritic structure. The dendritic tree is an arborization of total length *l*_*t*_ containing *B* generation of branches. For simplicity all branches of a generation *b* ∈ [1, *B*] have a length 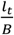. From one generation to the other, a branch divides into two branches where the diameter of the daughter branches follows Rall's 3/2 branching rule [17] 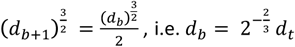 where *d*_*t*_ is the diameter of the root branch of the dendritic tree. Excitatory and inhibitory inputs are then spread homogeneously over the soma and dendritic tree according to the densities of synapses 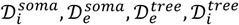. The parameters of the model are presented on Table 1.

### Model equations: synaptic input and passive properties

The cable equation describes the temporal evolution and spatial spread of the membrane potential along the branches of the dendritic tree [17]:

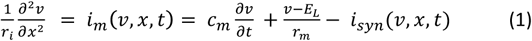

the membrane current *i*_*m*_(*ν*, *x*, *t*) is a linear density of current (the presented cable equation already includes the radial symmetry, i.e 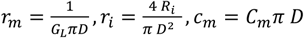, where *G*_*L*_, *R*_*i*_, *C*_*m*_ are the passive membrane parameters, see Table 1). Though the modeled system has several branches, the equation can be written as a single spatial dependency *x* because the symmetry of the model across branches imply that the properties of the input are identical at a given distance to the soma.

Synaptic input is modeled by local (infinitely small) and transient changes of membrane permeability to selective ionic channels. Both excitatory (accounting for AMPA synapses) and inhibitory synapses (accounting for GABAa synapses) are considered, their reversal potential is *E*_*e*_=0mV and *E*_*i*_=-80mV respectively. Each synaptic event is generated by a point process and its effect on the conductance is an increase of a quantity *Q*_*s∈*{*e*,*i*}_ followed by an exponential decay *τ*_*s∈*{*e*,*i*}_. The form of the synaptic current is therefore:

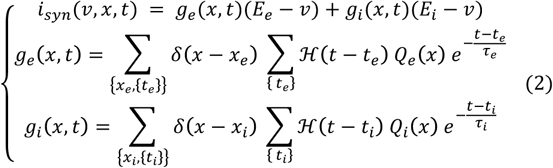

where *g*_*e*_ and *g*_*i*_ are linear densities of conductances. Each synapse, indexed by *S*, has a position *x*_*s*_ and a set of presynaptic events {*t*_*s*_}, hence the iteration over 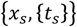 for the sum over synapses for each synaptic type. ℋ is the Heaviside step function. The presynaptic events {*t*_*s*_} are generated by point processes at fixed frequencies {*ν*_*s*_} with a given degree of synchrony, see details in the next section.

The model distinguishes two domains: a proximal domain (*x* ∈ [0, *l*_*p*_]) with the upper index *p* and a distal domain (*x* ∈ [ *l*_*p*_, *l*]) with the upper index *d* (see Figure **1**), where *l*_*p*_ is the length of the somatic compartment and *l* is the total length of the dendritic tree. The space-dependent quantities (presynaptic frequencies, synaptic quantal and synaptic decay time constant) can be written as:

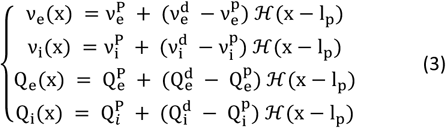

The continuity of the membrane potential and of the current at the boundaries between the proximal and distal part imply:

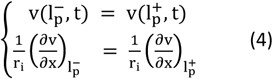

where the limit with upper index ± indicate the limit taken from the left or the right respectively.

At the soma, *x* = 0, we have a lumped impedance compartment. It has leaky RC circuit properties and also receives synaptic inhibition, the somatic membrane potential therefore follows:

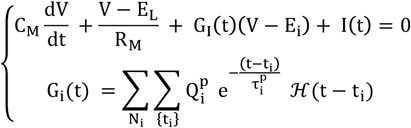

where *I*(*t*) is the time-dependent input current from the soma into the dendrite. *R*_*M*_ and *C*_*M*_ are the RC properties of the lumped compartment (capital letters will indicate the somatic properties throughout the calculus). *N*_*i*_ is the number of somatic synapses, each of them generates a point process {*t*_*i*_} of inhibitory synaptic events. The properties of the somatic synapses 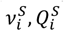 are equivalent to the proximal ones.

This equation with the membrane potential continuity will determine the boundary condition at the soma ( *x* = 0). We identify *V*(*t*) = ν(0, *t*), then *I*(*t*) is the current input into the dendritic tree at *x* = 0 so it verifies:

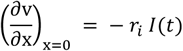

So:

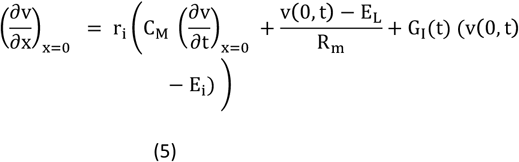

Finally, the last boundary condition is that all branches terminate with an infinite resistance that impede current flow (sealed-end boundary conditions):

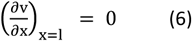

Together with the biased Poisson process for event generation (see the next section), the final set of equations that describes the model, is therefore the combination of Equations **1**, **2**, **3**, **4**, **5**, **6**.

### Model of presynaptic activity

Presynaptic activity is modelled as a discrete set of presynaptic events. Because of the apparent random spiking activity in the fluctuation-driven regime, the basis for the generation of those discrete events is the Poisson process. Nonetheless, we want to reproduce the additional degree of presynaptic synchrony found in neocortical assemblies that result mainly for two phenomena: 1) pairwise correlation between neurons and 2) multi-innervation of a cell by a presynaptic neuron. We therefore introduce a variable *S* that biases the event generation of the Poisson process (*S* ∈ [0,1]) by introducing coincident spikes. For simplicity in the analytical treatment, synchrony in presynaptic activity is not shared across different synapses. The relatively low range of synchrony values investigated here (*S* ∈ [0, 0.4]) also allows us to limit the number of coincident events to four events (quadruple events have a probability p≃ 0.06 for s=0.4, hence, even for the highest synchrony value, synchrony is dominated by pairs and triples of presynaptic spikes). Therefore, for a degree of synchrony *S*: single events have a probability 1 − *S*, double events have a probability *S* − *S*^2^, triple events have a probability *S*^2^ − *S*^3^ and quadruple events have a probability *S*^3^. To generate a biased Poisson process of frequency *ν* with a degree of synchrony *S*, we generate a Poisson process of frequency

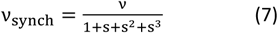

and we duplicate (from up to four events) each event according to their probabilities of occurrence.

This is a very simplistic and limited model of presynaptic synchrony but it is sufficient to reproduce the impact of synchrony on the quantities investigated in this paper (only the variance of the membrane potential fluctuations).

### Numerical implementation

The full model has been implemented numerically using the *NEURON* software [20]. The branched morphology was created and passive cable properties were introduced (see Table 1). The spatial discretization was *nseg*=30 segments per branch. On each segment, one excitatory and one inhibitory synapse were created, the shotnoise frequency was then scaled according to the segment area and the synaptic density to account for the number of synapses on this segment (using the properties of the Poisson process, N synapses at frequency *ν* is a synapse at frequency *N*_*ν*_. Custom event generation was implemented to introduce correlations (instead of classical *NetStim*) and fed *NetCon* objects attached to each synapses (*ExpSyn* synapses). Each simulation had a time step dt=0.01ms and a length of 10s, the simulation was repeated over 4 seeds to yield a mean and a standard deviation in the estimate of the membrane potential fluctuations at the soma (see Figure **3**B).

### Analytical derivation of the fluctuation properties: strategy

We present here a derivation that provides an analytical approximation for the properties of the fluctuations of the membrane potential at the soma for our model. Summing up its properties, we get: 1) a morphology with a lumped somatic compartment and a dendritic tree of symmetric branching following Rall's rule 2) conductance-based synapses 3) independent excitatory and inhibitory shotnoise input spread all over the morphology 4) asymmetric properties between a proximal part and a distal part and 5) a certain degree of synchrony in the pre-synaptic spikes.

The properties of the membrane potential fluctuations at the soma correspond to three stationary statistical properties of the fluctuations: their mean *μ*_*V*_, their standard deviation *σ*_*V*_ and their *global* autocorrelation time *τ*_*V*_. Following [6], we emphasize that the *global* autocorrelation time is a partial description of the autocorrelation function (as the autocorrelation function is not exponential) but it constitutes the first order description of the temporal dynamics of the fluctuations.

A commonly adopted strategy in the *fluctuation-driven* regime to obtain statistical properties is to use stochastic calculus after having performed the *diffusion approximation*, i.e. approximating the synaptic conductance time course by a stochastic process [33]. This approach is nonetheless not easily generalizable to conductance input in an extended structure and render the inclusion of asymmetric properties (proximal vs distal) complicated. We rather propose here an approach that combines simplifying assumptions and analytical results from point process theory, it extends the approach proposed in Kuhn et al. [19] to dendritic structures following Rall's branching rule. For each set of synaptic stimulation 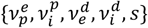, the derivation corresponds to the following steps:

I. We transform the dendritic structure to its equivalent cylinder. The reduction to the equivalent cylinder is "activity-dependent" and captures the changes in membrane properties that results from the mean synaptic conductance levels.
II. We derive a mean membrane potential *μ*_*V*_(*x*) corresponding to the stationary response to constant densities of conductances given by the means of the synaptic stimulation. We use this space-dependent membrane potential *μ*_*V*_(*x*) to fix the driving force all along the membrane for all synapses. The relation between synaptic events and the membrane potential now becomes linear.
III. We derive a new cable equation that describes the variations of the membrane potential around this *μ*_*V*_(*x*) solution.
IV. We calculate the effect of one synaptic event on a branch *b*, *b* ∈ [1, *B*] at a distance *x*. We calculate the post-synaptic membrane potential event PSP_*b*_(*x*, *t*) at the soma resulting from *b* synchronous synaptic events occurring at the distance *x* from the soma. We approximate the effect of only one event by rescaling the response by the number of input 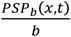.
V. We use shotnoise theory to compute the power spectrum density of the membrane potential fluctuations resulting from all excitatory and inhibitory synaptic events (including the synchrony between events).

The full derivation has been conducted with the help of the python modulus for symbolic computation: *sympy*. The resulting expression were then exported to *numpy* functions for numerical evaluation. Details are provided in **Supp. material**

### From fluctuation properties to spiking probability

How layer V pyramidal neurons translate membrane potential fluctuations into a firing rate response was the focus of our previous communication [6]. We re-use here the same dataset, i.e. the individual characterizations over n=30 single neurons of the firing responses 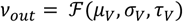.

Importantly, our study introduced four quantities to describe the mean properties of a single neuron response in the fluctuation-driven regime: (i) a measure of the excitability given by the mean phenomenological threshold 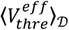 of the firing response function, (ii) a sensitivity to the mean of the fluctuations 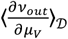, quantifying how much a mean depolarization is translated into a change in firing rate, (iii) a sensitivity to the amplitude of the fluctuations 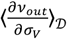, quantifying how much an increase of the standard deviation of the fluctuations is translated into a change in firing rate, and (iv) a sensitivity to the speed of the fluctuations 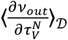, quantifying how much a change in the autocorrelation time of the fluctuations is translated into a change in firing rate. Note that in the main text, we removed the n=3/30 cells that had too low excitabilities, as it is hard to conduct an analysis on 6 orders of magnitudes, *ν* ∈ [10^−5^, 10^1^] Hz (for data visualization). This reduction limited the output variations to *ν* ∈ [10^−2^, 10^1^] Hz. In the supplementary figure S3, we reintroduce the discarded cells and we show that they do not affect the results presented in the main text.

### Experimental preparation and electrophysiological recordings

Experimental methods were identical to those presented in [6]. Briefly, we performed intracellular recordings in the current-clamp mode using the perforated patch technique on layer V pyramidal neurons of coronal slices of juvenile mice primary visual cortex. For the n=13 cells presented in this study, the access resistance *R*_*S*_ was 13.3MΩ ± 5.4, the leak current at -75mV was -25.7pA±17.3, cells had an input resistance *R*_*m*_ of 387.3M Ω ± 197.2 and a membrane time constant at rest of 32.4ms±23.1.

### Input impedance characterization

To determine the input impedance at the soma, we injected sinusoidal currents in the current-clamp mode of the amplifier (Multiclamp 700B, Molecular Devices), we recorded the membrane potential response to a current input of the form *I t* = *I sin*(2*πft*), we varied the frequencies *f* and amplitudes *I* over 40 episodes per cell. The frequency range scanned was [0.1, 500] Hz. For each cell, we determined manually the current amplitude *I*_0_ that gave a ∼5mV amplitude in a current step protocol, from this value, the value of *I* was scaled exponentially between *I*_0_ at 0.1 Hz and 50 *I*_0_at 500Hz. The reason for varying the current amplitude (and not only the oscillation frequency) in those input impedance protocols is to anticipate for the low pass filtering of the membrane and insure that the membrane potential response at high frequencies is far above the electronic noise level ∼ 0.1 mV.

After removing the first 3 periods of the oscillations (to avoid transient effects), we fitted the membrane potential response to the form: *V*(*t*) = *E*_*L*_ + *R I sin*(2*πft* − ϕ), where *E*_*L*_, *R*, *ϕ* were fitted with a least-square minimization procedure. The frequency dependent values of *R* and *ϕ* give the modulus and phase shift of the input impedance presented in Figure **3**A.

### Fitting passive properties and a mean morphology

Because the variables combined discrete (the branch number) and continuous variables, the minimization consisted in taking the minimum over a grid of parameters. The parameter space had 7 dimensions: the branch number 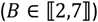, the somatic length ( *l*_*s*_ ∈ [5, 20]*μ* m), the total length of the tree ( *l*_*t*_ ∈ 300, 800 *μ* m), the diameter of the root branch ( *d*_*t*_ ∈ 0.5, 4 *μ* m), the leak specific resistance ( *r*_*m*_ ∈ 100, 1000]*μ* S/cm^2^), the intracellular resistivity ( *r*_*i*_ ∈ [10, 90] Ω .cm), the specific capacitance ( *c*_*m*_ ∈ [0.8, 1.8] *μ*F/cm^2^). Each dimension was discretized in 5 points, the scan of the 7 dimensional space then consisted in finding the least square residual of the product of the modulus and phase of the impedance over this 5^7^ points. The resulting parameters are shown on Table 1.

## Acknowledgments

We would like to thank Gilles Ouanounou for his help during electrophysiological recordings and Gérard Sadoc for assistance with the acquisition software. We also thank Manon Richard and Aurélie Daret for animal facilities. Research supported by the CNRS and the European Community (Human Brain Project, H2020-720270).

## Supporting Information

**Figure S1.**
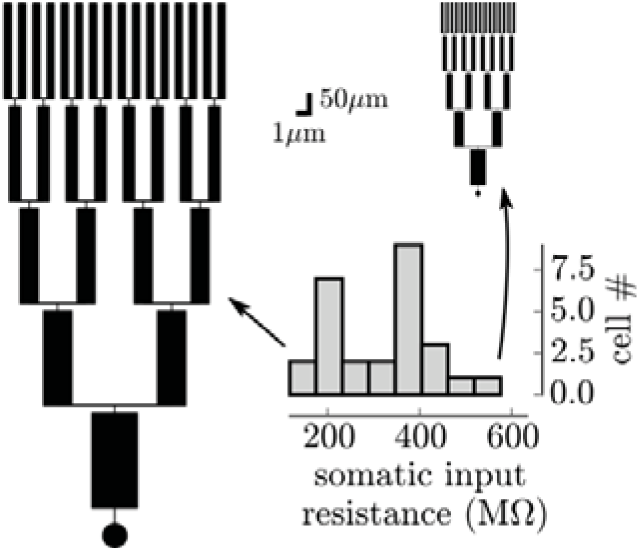
Heterogeneity in morphologies. Graphical representation of the estimated morphologies of the largest (the lowest input resistance) and the smallest (the highest input resistance) cells present in the dataset.

**Figure S2.**
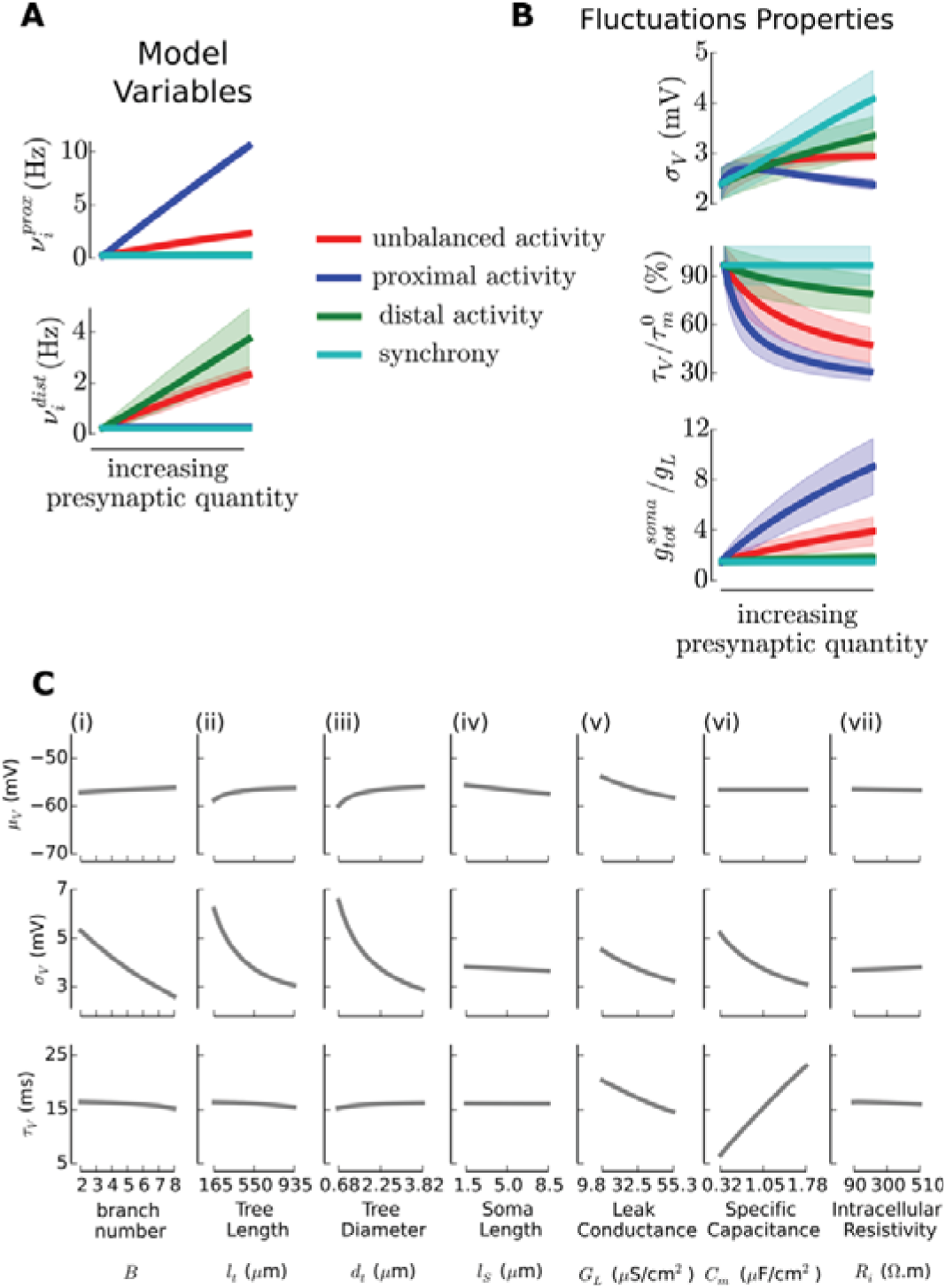
Variability in the fluctuations properties introduced by different morphologies. **(A)** Variability in the model variables across protocols introduced by the different morphologies (sizes). We show only the quantities that vary across cells, all other variables 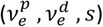 are fixed across cells. The balance *μ*_*V*_ is adjusted for each cells and the cells have different surfaces, so different number of synapses (and especially different ratio of excitatory to inhibitory numbers) hence the need to adjust inhibitory activity slightly differently for each cell. **(B)** Variability in the properties of the membrane potential fluctuations (standard deviation *σ*_*V*_ and autocorrelation time *τ*_*V*_) across protocols introduced by the different morphologies (sizes). We show only the quantities that vary across cells, *μ*_*V*_ is fixed across cells by design. **(C)** Dependency of the fluctuations properties as a function of the morphology parameters (all parameters are varied of -70% and +70% around those of the mean model, one parameter is varied while all others are fixed to those of the mean model, see Table 1). We fixed the presynaptic stimulation to a level of 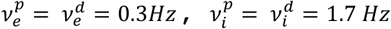 and *S* = 0.05. Globally, the mean polarization *μ*_*V*_ is poorly affected by the morphology, given the balanced nature of the input and the homogenous spread of synapses on the membrane. On the other hand, the morphology parameters strongly affect the properties of the amplitude and speed of the fluctuations. **(i)** Dependency on the branching number. Because the branching number has an impact on the area, the number of synapses increases with the number of branches, then the amplitude *σ*_*V*_ of the fluctuations strongly decreases with the number of branches because of the law of large numbers. **(ii)** Dependency on the tree length. The same argument as in (i) holds. The length of the tree increases the number of synapses so that it reduces the amplitude of the fluctuations. **(iii)** Dependency on the tree diameter. The same argument as in (i) holds again. The diameter of the tree increases the number of synapses so that it reduces the amplitude of the fluctuations. **(iv)** Dependency on the size of the soma. Here, this increases only the number of inhibitory synapses at the soma. Within this range this has very weak effects on the fluctuations. **(v)** Dependency on the leak conductance. The leak conductance sets the amplitude and low pass filtering of individual post-synaptic events (because *τ*_*m*_ ∼ *C*_*m*_/*G*_*L*_, see correction introduced by the ongoing activity in the main text). An increasing leak conductance will “shunt” post-synaptic events. This lowers the amplitude of the fluctuations *σ*_*V*_ and lowers the autocorrelation time of the fluctuations *τ*_*V*_. **(vi)** Dependency on the specific capacitance. Again because *τ*_*m*_ ∼ *C*_*m*_/*G*_*L*_, an increasing capacitance renders the fluctuations much slower (increasing *τ*_*V*_). Also because PSP events have a much lower frequency content, they lead to a reduced amplitude of the fluctuations *σ*_*V*_. **(vii)** Dependency on the intracellular resistivity. This has a moderate effect within this range.

**Figure S3.**
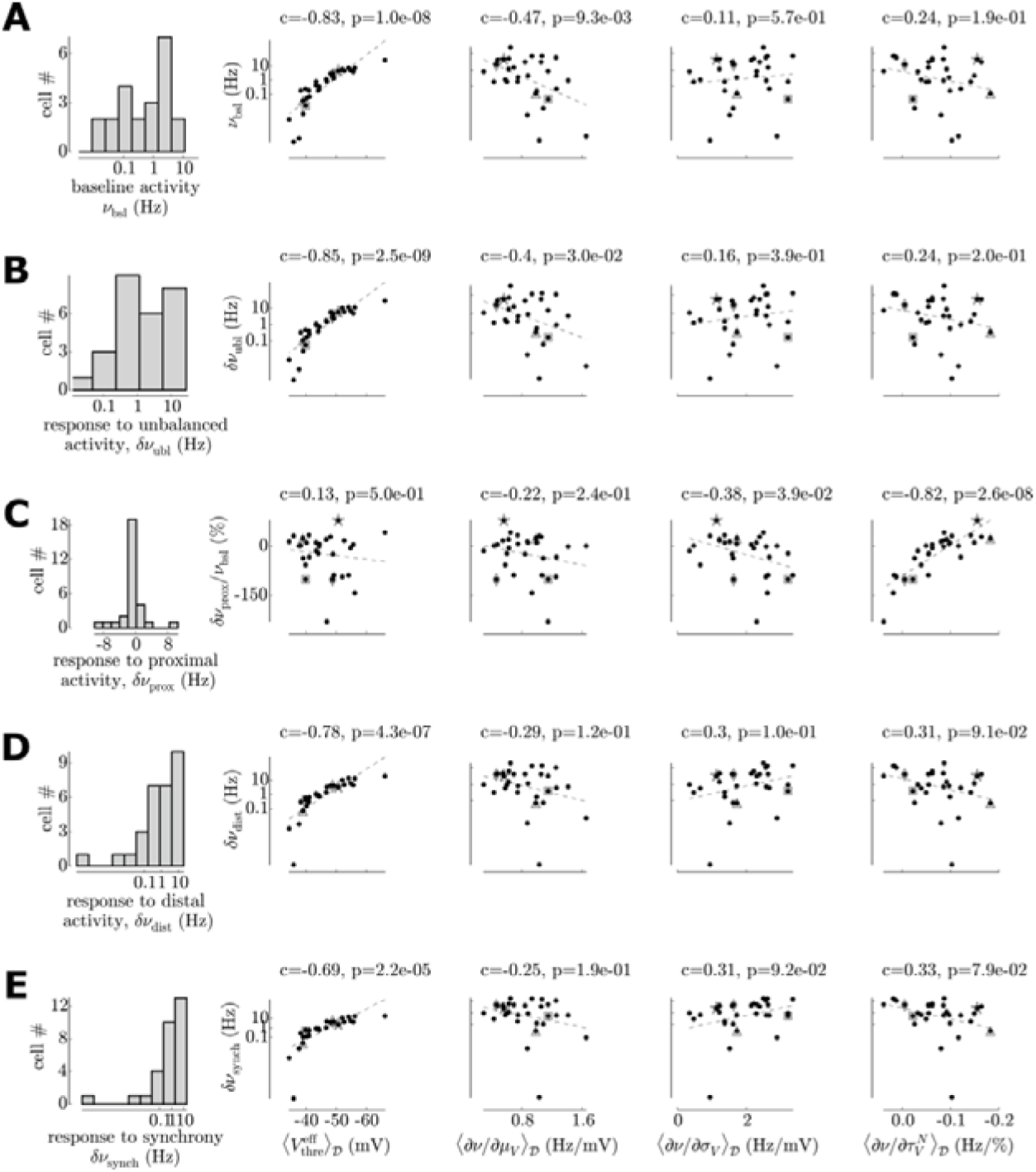
Re-including all cells in the study of the correlation between the firing rate responses and the firing response function characteristics. We re-introduce here the recorded cells of very low excitability that produce very low firing rate responses (*ν*_*out*_ < 10^−3^) and partially impede data visualization (because they shift extend the axis limits far from the majority of the data). Note that the strong correlations discussed in the result are not affected.

**Figure S4.**
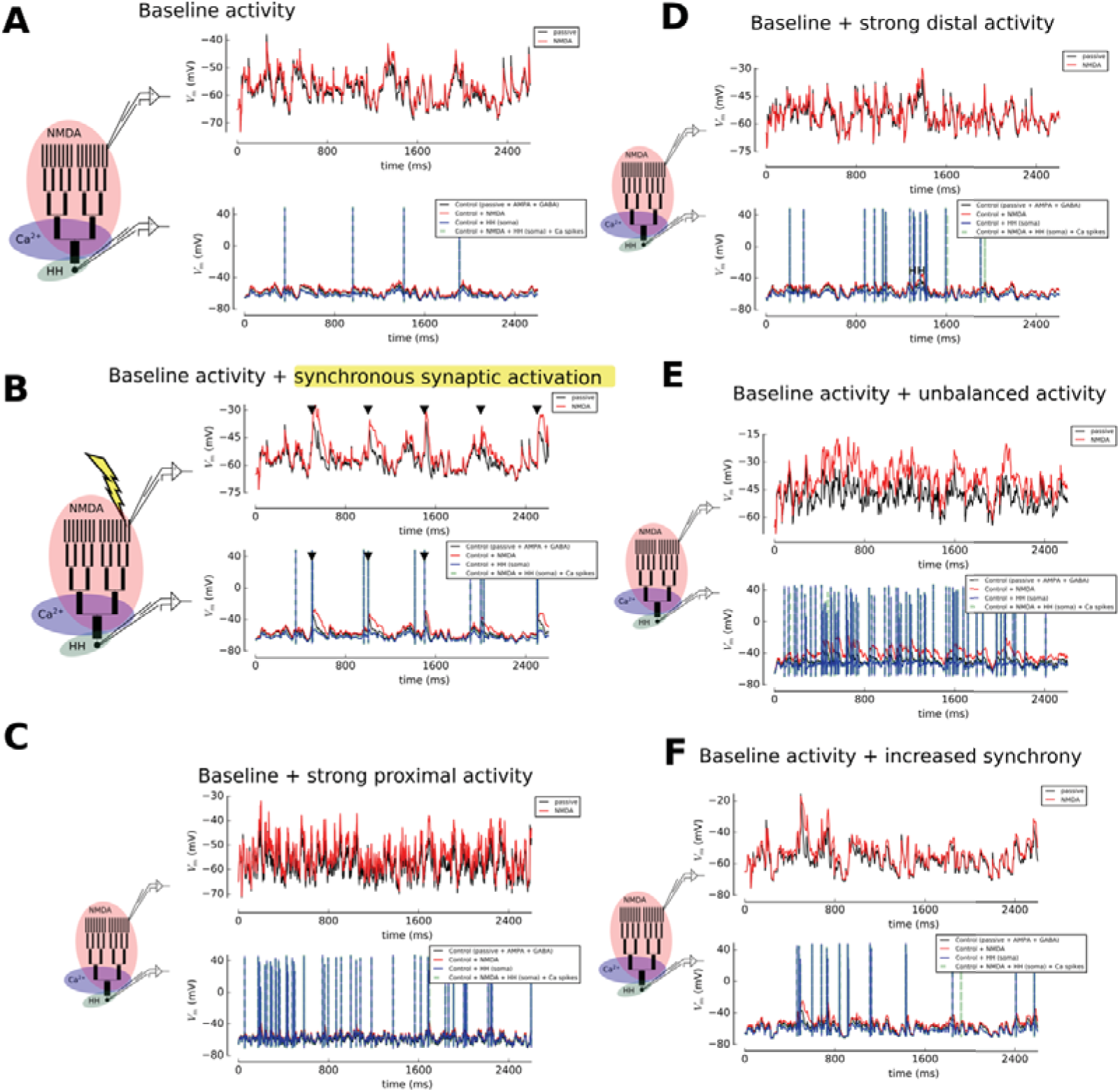
Including dendritic non-linearities (numerically) within our framework for single cell computation. Active mechanisms are taken from Larkum et al. [31]. On top of each panel, we present the recording of the membrane potential in the far distal dendrite (depicted with the upper electrode on the drawing) in the Control and Control+NMDA case. On the bottom, we present the membrane potential recorded at the soma (depicted with the lower electrode on the drawing) in four cases, the Control, the Control+NMDA case, the Control case with a HH model inserted in the soma and the Control+NMDA+Ca^2+^spikes+HH(soma). **(A)**Response to baseline activity. 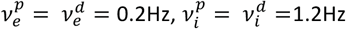 and *S*=0.05. **(B)** Response to baseline activity with the synchronous activation of 20 synapses every 500ms (marked by a triangle), note the impact of the NMDA mechanism under this stimulation type. **(C)** Response to an increase of proximal activity with respect to the baseline level 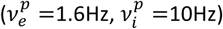. **(D)** Response to an increase in distally targeting presynaptic activity 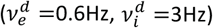. **(E)** Response to an increase in unbalanced activity 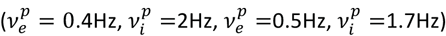. **(F)** Response to an increase in presynaptic synchrony (*S* =0.4).

**Figure S5.**
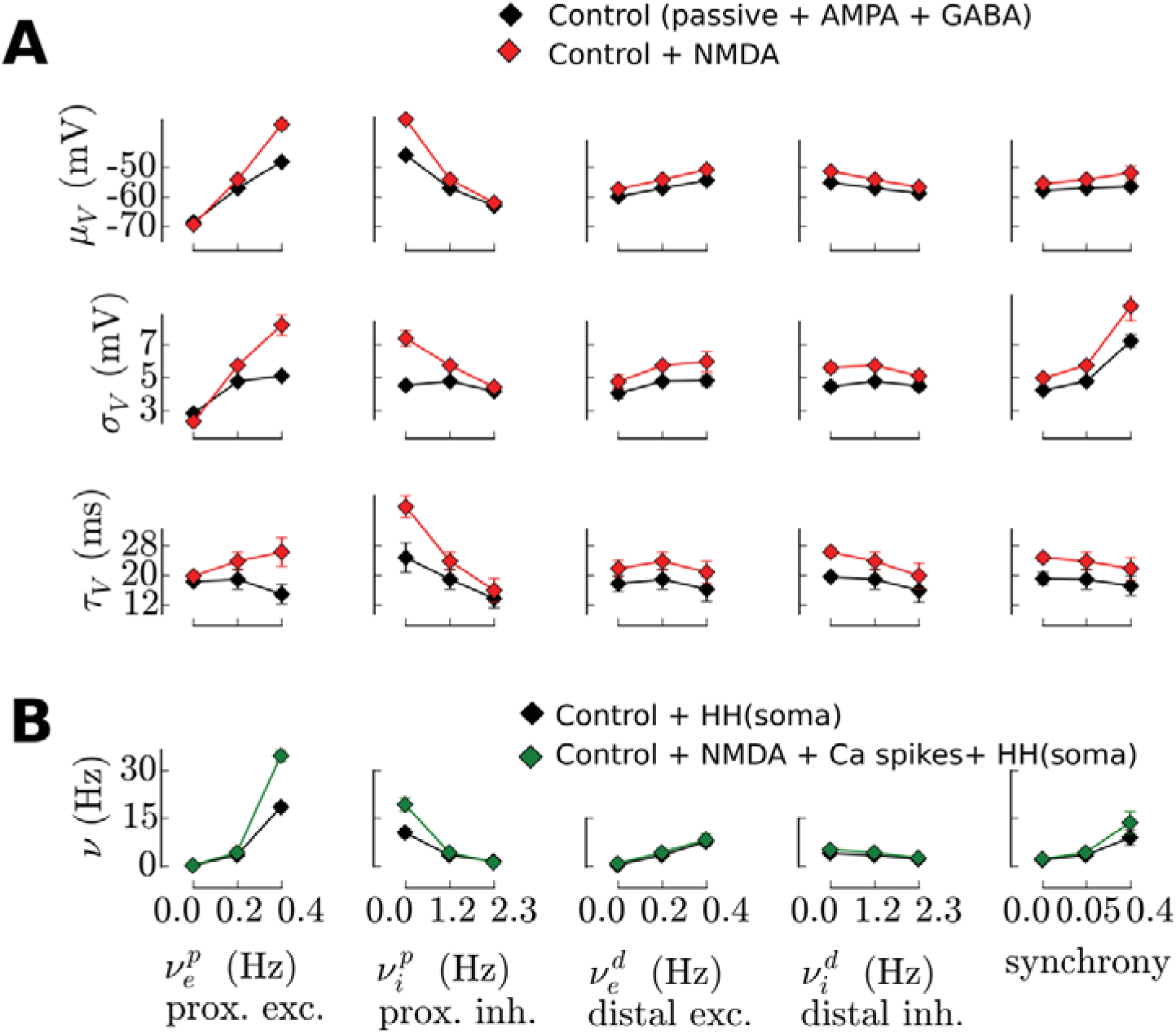
Active dendritic mechanisms do not qualitatively affect the relationship between presynaptic activity and firing rate provided that excitation does not unbalance inhibition (*μ*_*V*_ ≲ −50mV), and provided that synchrony remains in a relatively low range (*s* ≲ 0.4). Because our framework for single cell computation consists in (i) evaluating analytically the somatic subthreshold membrane potential fluctuations at the soma as a function of the presynaptic quantities and (ii) convert those fluctuations into a spiking probability thanks to a *firing response* function determined *in vitro* in individual neuron, it needs to be tested against two phenomena: (I) whether active dendritic mechanisms qualitatively affect the relationship between presynaptic quantities and somatic membrane potential fluctuations, i.e. checking the validity of step (i), and (II) whether dendritic non-linearities introduce a coupling between presynaptic activity and spike emission that renders the decomposition into the successive steps i & ii problematic. Those phenomena will naturally appear to some extent and will produce quantitative deviations with respect to our predictions (that do not include dendritic non-linearities, e.g. trivially, all cells should be more excitable). As our results do not rely on the absolute values of the input-output functions but rather on the relative behaviors from cell to cell (that emerge from differences across individual firing response functions), we only look for putative “qualitative deviations”. **(A)** We first investigate whether active dendritic mechanisms qualitatively affect the relationship between presynaptic quantities and somatic membrane potential fluctuations. We observe that NMDA channels exert an additional small depolarization (increased *μ*_*V*_), they also increase the standard deviation of the fluctuations because they increase the amplitude of depolarizing events (increased *σ*_*V*_) and they slow down the fluctuations because they increase the low frequency content of post-synaptic depolarization (increased *τ*_*V*_). We also observe, that, despite this small shift in all quantities, the qualitative trend of all curves remains identical. Only for high excitation, a qualitative difference can be seen (see bottom left plot): increasing excitation makes *V*_*m*_ fluctuations faster in the passive case, whereas increasing excitation makes fluctuations slower in the presence of NMDA currents (decreasing vs increasing *τ*_*V*_ curve respectively). **(B)** We then investigate whether dendritic non-linearities introduce a coupling between presynaptic activity and spike emission that would complicate our picture. This was tested by including both NMDA and Ca^2+^ currents. We first included numerically a spike emission model at the soma (Hodgkin-Huxley like, Na+ and K+ currents) in the morphological model, to allow the model to produce the full input-output function (from presynaptic quantities to spiking probability). Trivially, we observe an increased spiking probability because cells are more excitable (NMDA and Ca^2+^ currents are depolarizing currents). Notably, when excitation does not balance inhibition (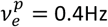 or 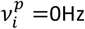), the spiking probability is greatly enhanced (almost doubled). For the other variables, those effects are much more moderate.

